# Gut meta-virome reveals potential zoonotic pathogens and environmental RNA viruses carried by non-human primates

**DOI:** 10.1101/2025.01.07.631708

**Authors:** Yujie Yan, Yuhang Li, Linshan Yang, Haojie Wu, Fan Wu, Hongli Chang, Zhengfeng Hu, Shujun He, Yi Ren, Lifeng Zhu, Baoguo Li, Songtao Guo

## Abstract

The majority of human infectious diseases originate from mammals and are inherently zoonotic. Non-human primates (NHPs) are not only carriers of many zoonotic pathogens, but also the best intermediary for virus shift from harmless to harmful due to their similar phylogenetic relationship with humans. Knowledge of NHP viral composition and its underlying information can therefore provide an assessment of the risk of cross-species transmission and spillover of zoonotic diseases. However, studies that have successfully eliminated the effects of different natural habitat environments on viral carriage have been limited, making it difficult to identify which zoonotic viruses and potentially species-barrier-crossing viruses are carried by widely distributed NHP hosts. Here, we analyze viruses excreted in feces from NHP hosts and reveal the presence of possible zoonotic pathogens. Most of the viruses found come from plants that are eaten by them. Analysis of the evolutionary relationships of potential pathogens suggests that other clinical isolates are genetically correlated to HIV, PBV, and EV we identified. This study provides foundational data for surveillance of NHP enteroviruses that could help to be challenging to track potential human viruses carried by NHP in the absence of clinical surveillance.

**Importance:** As a result of internationalization and other factors, many viruses have broken through the limits of geographical units and spread into human society, causing many emerging and re-emerging infectious disease hazards. However, the virus spreads and is by no means infectious to humans overnight. There is a phenomenon of host shift, in which the virus does not necessarily cause disease in an intermediate host, but causes significant damage when transmitted to humans. We focus on captive non-human primates, close relatives of humans in cities, because the phenomenon of host shifts is related to both biological similarity and geographical proximity of the host. Primates may carry zoonotic viruses and viruses that are currently incapable of infecting humans but have evolved with pathogenic potential. In summary, comprehensive viral surveillance and screening of captive nonhuman primates is necessary.

## 1. Introduction

Globalization and human activities have led to rapid spread of viral zoonoses pathogens across geographical units. More than 60% of known human infectious diseases and more than 70% of emerging infectious diseases are transmitted from vertebrates to humans (1, 2). Some zoonotic diseases caused by underlying viruses have not yet been fully explained, partially due to the fact that human knowledge and mastery of animal viruses is still largely inadequate. For example, the causes of the vast majority of human diarrhea cases remain unknown (3), and unidentified animal derived pathogens may be present.

It has recently been discovered that the infectious ability of viruses to move from animal to human and from harmless to harmful does not happen overnight. The barrier between humans and animals is broken by the evolution and mutation of viruses that use animals as transit hosts -- a phenomenon also known as ‘host shifts’ (4, 5). Many pathogens that have successfully crossed the species barrier may not be pathogenic or exhibit clinical symptoms in natural and/or intermediate hosts. This makes some viruses that derive from animal hosts and are evolutionarily potentially harmful to humans often neglected (6). An axample is *Simian immunodeficiency virus* (SIV), which may have gone unsupervised and unnoticed because it does not satisfy Koch postulates and does not necessarily lead to morbidity in simian hosts (7, 8), leading to missed prevention of HIV in humans. Therefore, comprehensive viral surveillance of animals is essential. The application of high-throughput sequencing technology for comprehensive viral surveillance mapping at the human-animal-ecosystem interface can provide opportunities for the discovery of both known and as-yet-unseen pathogens (9–12). This helps us to determine the viral composition of animals, as well as to assess the evolutionary processes and potential pathogenicity of viruses.

Ongoing scientific surveillance suggests that viral shift is more likely to occur between new hosts of relevance and old hosts (e.g., with similar immune responses or life history characteristics) (13–15). Non-human primates (NHP) are thus the most critical host transit for viruses to escape evolutionary dead ends and mutate into zoonoses. Apart from the familiar HIV, major infectious diseases shifte<u>d</u> from the NHP to humans include the *Simian foamy virus* (SFV), *Yellow fever virus* (YFV), and *Zika virus* (ZIKV), among others (16–18). In addition, recent studies have identified a variety of known or novel viruses with zoonotic potential, such as *Coxsackieviruses* (19), *Enteroviruses* (20, 21), and *Picobinaviruses* (21, 22), in the gut of both wild and captive NHP. These NHP viruses will shift from primates to humans and evolve to pose a threat to human health. However, human knowledge of NHP virus diversity, host range, phylogeny and cross-species transmission factors remains fragmented and inadequate. This constrains early warning, prevention and control of emerging and re-emerging zoonotic infections.

A assumption is that in order to track and identify potential human viruses that may arise, we should focus on the NHPs that are kept in contact with people in cities. The complexity and diversity of habitat types of wild NHP (23), with sufficient potential to carry important pathogens related to geographic or environmental factors, has led to the neglect of a number of viruses that are currently unable to infect humans but have the evolutionary potential to cause disease. Moreover, host shifts of viruses are associated with geographic proximity between hosts (4). Of the NHP pathogens for which a primary route of transmission could be identified, 45% could be transmitted by close non-sexual contact, 43% by non-intimate contact, and 32% by arthropod vectors (a combined pattern of these routes of transmission exists) (24). This gives more opportunities for the shifting of potential zoonotic pathogens from animals kept in cities. For example, a case of monkey B virus transmission from a captive monkey to a human was reported in Beijing, China, in 2021, resulting in a human death (25). After reviewing global primate virus infections and transmission, Liu et al. called for comprehensive viral surveillance of captive NHPs to identify viruses with zoonotic potential, especially in old world monkeys and monkeys in regular contact with humans (22). However, Patouillat et al. reviewed 152 studies on zoonotic viruses and found that only a few NHP species had been oversampled (notably species of the genus *Macaca*), while many others had been ignored (26).

To predict potential zoonotic viruses transmitted from NHP to humans, here we conducted food recording and comprehensive screening of RNA meta-virome targeting 15 primate species with different natural habitats and uniformly raised in a zoo of South China. These NHPs include prosimians, new world monkeys, old world monkeys, and apes. Our aims were (i) the ability to study gut viruses carried by captive individuals in unprecedented detail across multiple NHP species, and (ii) the mining of known or potential zoonotic pathogens. This study emphasizes the importance of NHP virus research and provides insights into the risk of viral shift in species without adequate clinical surveillance.

## 2. Results

### 2.1 Non-human primates investigated

We investigated gut viruses in captive NHPs at a zoo in southern China and recorded observations of the corresponding diets (Table S1). Our animal samples were collected before January 2020 and were not disturbed by the SARS-CoV-2 pandemic. And, despite differences in natural habitats (Figure 1a), our NHP fecal samples were collected in the same area under captive conditions, eliminating the influence of differences in environmental factors on viral infections in animals. None of the individual animals took or injected drugs during our observation and sample collection period, and there were no obvious signs or symptoms of illness. The full collection of 45 fecal samples consisted of 15 individual animals from 15 species, representing five NHP families and four evolutionary taxa (Figure 1b). Subsequently, fecal samples were pooled according to species and divided into 15 pools for meta-virome sequencing.

**Figure 1.**
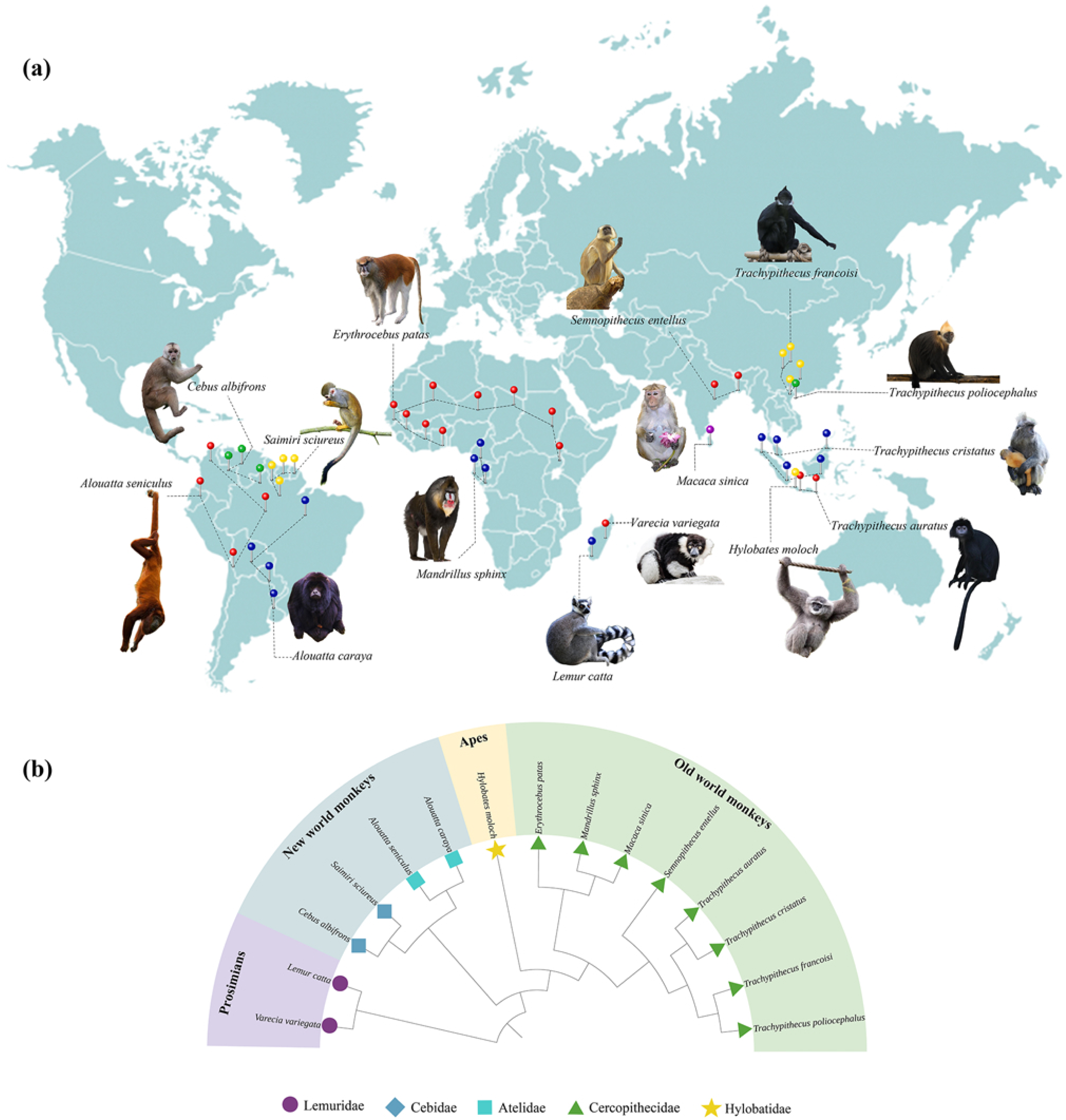
General information on 15 samples. (a) natural habitats of non-human primate species; (b) host species evolutionary relationships

### 2.2 General information of virome sequencing and assembly

After Illumina sequenced the viruses in the fecal samples, a total of 721,008,076 raw reads were obtained. A total of 261,089,182 clean reads were reassembled into 168,721 contigs over 300bp in length based on the comparison of reference data. Virus reads per sample ranged from 27,009-25,370,234 ( average of 7,328,217 reads), with 45.3%-100% of the reads representing RNA viruses (median 98.88%). Ultimately, 3,188 vOTUs longer than 300 bp were identified in the total sample virome (Table 1).

**Table 1.** General information of sequenced data.

| Sample | Raw Reads | Filtered reads | Virus reads | % of RNA virus | N50 | No. of contigs | Longest contig | N50/De novo | No. of contigs/De novo | Longest contig/De novo | Virus spices |
| --- | --- | --- | --- | --- | --- | --- | --- | --- | --- | --- | --- |
| *Varecia variegata* | 34,218,384 | 7,965,457 | 27,009 | 98.88 | 512 | 2,526 | 23,319 | 737 | 121 | 3564 | 16 |
| *Lemur Catta* | 44,296,225 | 27,353,850 | 21,685,504 | 100 | 534 | 3,833 | 35,552 | 602 | 231 | 6,886 | 24 |
| *Cebus albifrons* | 53,660,046 | 14,442,291 | 5,745,719 | 99.99 | 556 | 2,859 | 35,611 | 605 | 571 | 6,520 | 26 |
| *Saimiri sciureus* | 41,842,785 | 12,025,134 | 5,445,104 | 45.3 | 536 | 5,659 | 8,746 | 682 | 374 | 5,855 | 49 |
| *Alouatta caraya* | 48,210,054 | 13,207,522 | 30,539 | 76.73 | 653 | 12,634 | 72,036 | 2,348 | 43 | 4,001 | 10 |
| *Alouatta seniculus* | 46,267,188 | 10,452,819 | 96,824 | 81.31 | 543 | 16,336 | 12,225 | 812 | 125 | 9,261 | 35 |
| *Mandrillus sphinx* | 54,635,752 | 21,081,545 | 208,743 | 52.51 | 705 | 40,230 | 45,238 | 660 | 368 | 6,217 | 57 |
| *Macaca sinica* | 55,063,088 | 26,989,655 | 170,137 | 50.69 | 644 | 11,581 | 14,297 | 982 | 136 | 6,394 | 26 |
| *Erythrocebus patas* | 57,850,019 | 11,167,535 | 160,700 | 96.58 | 647 | 18,416 | 87,372 | 1,002 | 195 | 7,066 | 32 |
| *Trachypithecus francoisi* | 43,139,514 | 8,274,713 | 351,507 | 99.4 | 559 | 12,794 | 36,714 | 1,055 | 136 | 5,271 | 22 |
| *Trachypithecus poliocephalus* | 48,760,318 | 29,384,163 | 24,869,974 | 99.99 | 588 | 10,983 | 43,401 | 464 | 301 | 5,322 | 17 |
| *Trachypithecus cristatus* | 45,885,998 | 29,332,394 | 25,370,234 | 99.99 | 740 | 3,021 | 72,522 | 457 | 240 | 5,887 | 12 |
| *Semnopithecus entellus* | 44,471,480 | 9,349,489 | 1,069,576 | 99.69 | 620 | 17,781 | 69,902 | 767 | 143 | 5,348 | 25 |
| *Trachypithecus auratus* | 56,374,609 | 30,385,391 | 24,662,896 | 100 | 610 | 3,644 | 35,659 | 406 | 157 | 2,400 | 7 |
| *Hylobates moloch* | 46,332,616 | 9,677,224 | 28,783 | 87.8 | 609 | 6,424 | 99,760 | 2,303 | 47 | 5,973 | 18 |

### 2.3 Characterization of viral communities

Sequence annotations showed that a total of 376 viral species were found in the 15 samples, and the number in each sample is shown in Table 1. Only two viruses were shared across the 15 samples, *Hibiscus latent Fort Pierce virus* (HLFPV) and *Hibiscus latent Singapore virus* (HLSV). The upsets show that mandrill (*Mandrillus sphinx*) samples had the most unique virus spices, followed by squirrel monkeys (*Saimiri sciureus*) (Figure 2a). Most of the virus sequences come from viruses that infect plants, followed by animal viruses that can infect a variety of insects or vertebrates (Figure 2b).

**Figure 2.**
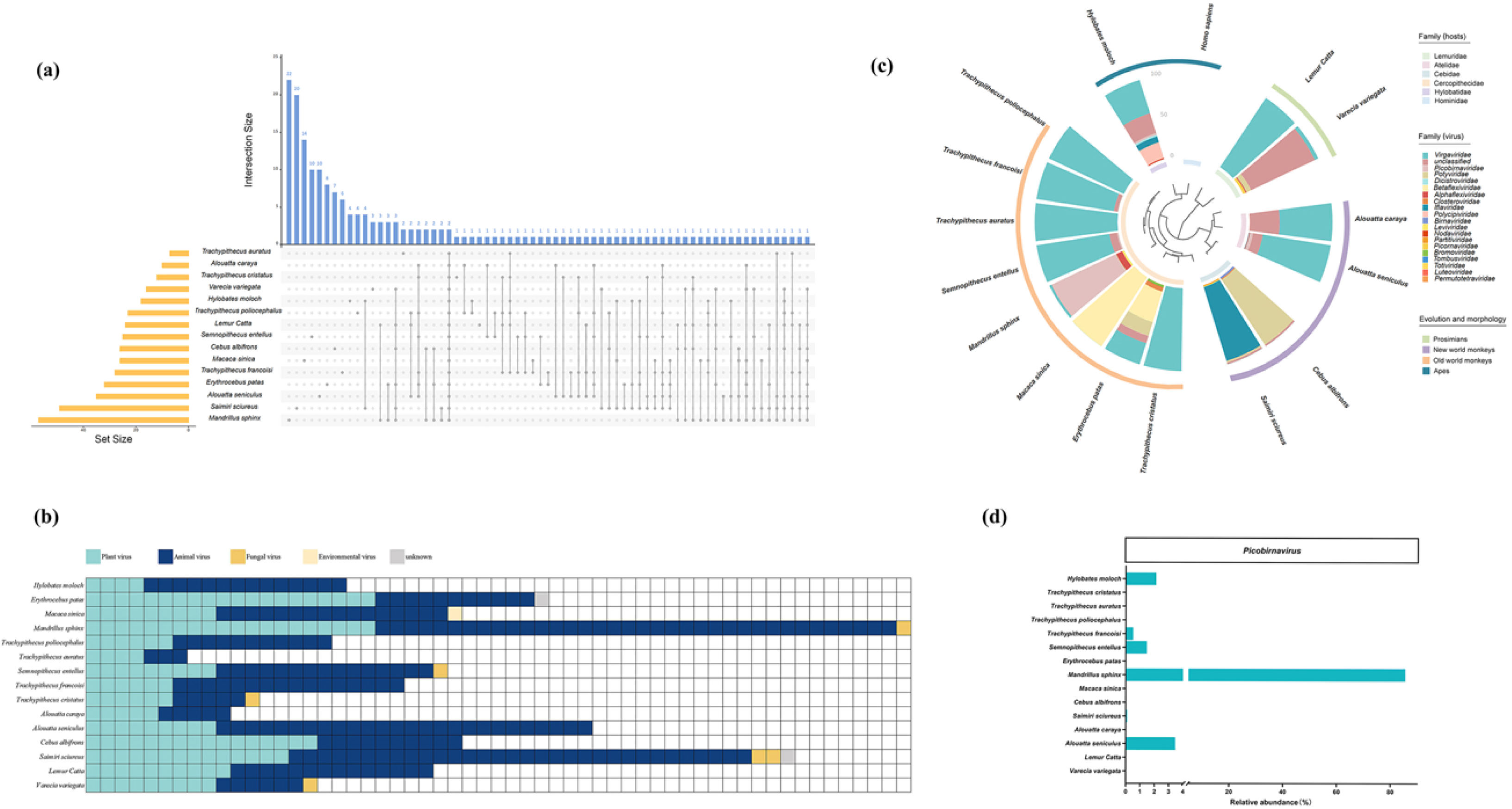
General characteristics of the viral community. (a) overlap of RNA virus species in different samples; (b) statistics on the type of RNA virus-derived sources in different samples; (c) distribution of abundance at the RNA virus family level and evolutionary information on hosts; (d) abundance of important Picobirna viruses

A total of 22 families were distributed among all samples in the classified virus taxa. Except for the unclassified families, the most abundant and widely distributed of them were *Virgaviridae* and *Picobirnaviridae* (Figure 2c). Despite differences in abundance at the family level among the 15 samples, viruses of the *Virgaviridae* had an extremely high percentage of abundance in four species of langurs of the genus *Trachypithecus* as well as in ring-tailed lemur (*Lemur catta*). Except for the francois’ langur (*Trachypithecus francoisi*), which accounted for 94.67%, the other four hosts accounted for more than 99.9%. The *Picobirnaviridae*, which are transmitted in mammals, were widely present in 13 samples, especially in the mandrill sample with an abundance accounting for more than 85.48% (Figure 2d).

The heatmap also showed that the relative abundance of viruses in the 15 samples differed significantly at the family level and genus level (Figures 3a, 3b). Unclassified families are significantly more abundant in black-and-white ruffed lemur (*Varecia variegata*) than in other species, while unclassified genera dominate in black-and-white ruffed lemur and squirrel monkey.

**Figure 3.**
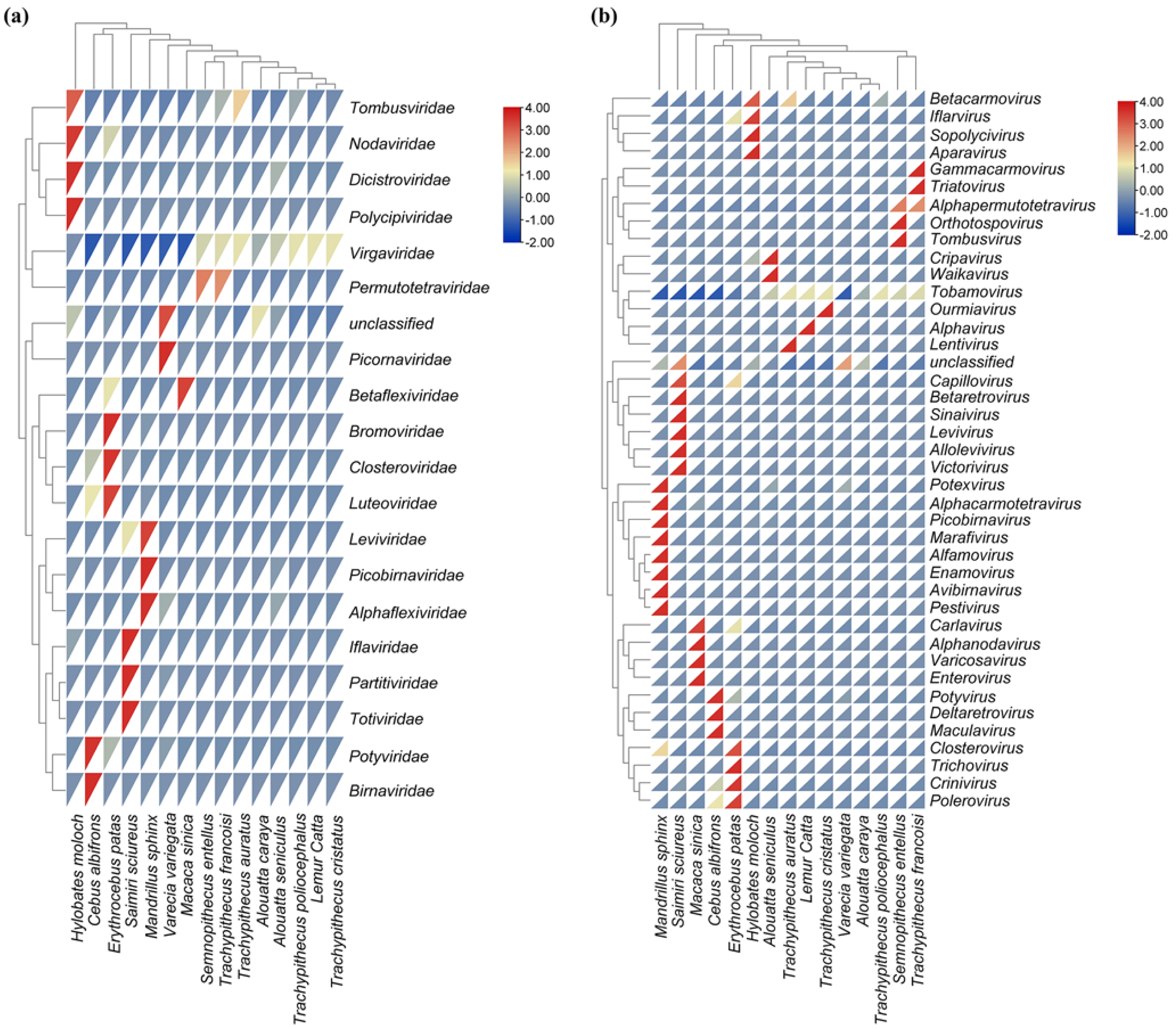
Viral community abundance characteristics. (a) Differences in relative abundance at the family level for 15 samples; (b) differences in relative abundance at the genus level for 15 samples

### 2.4 Detected viruses potentially infecting vertebrates

Potential pathogens with 33 predominantly vertebrate infections were identified in 15 primate fecal samples (Table 2). *Picobirnavirus* dominated all the potential pathogenic viruses identified, with all types of *Picobirnavirus* accounting for 6.2% of the total abundance of identified viruses. Also detected were the zoonotic enteroviruses *Enterovirus A*, *WUHARV Enterovirus2*, *WUHARV Enterovirus3*; the mosquito-borne *Guadeloupe Culex tymo-like virus*, *Mosinovirus*, *Culex originated Tymoviridae-like virus*; *Arenavirus*, *Astrovirus VA4* that may cause hemorrhagic fever; *Human blood-associated dicistrovirus*, *Semliki Forest virus*. In addition, low abundance of human pathogens were found, such as *human immunodeficiency virus 1* (HIV-1), where only one sequence was detected in a sample of Javan langur (RPKM = 0.5385).

**Table 2.**
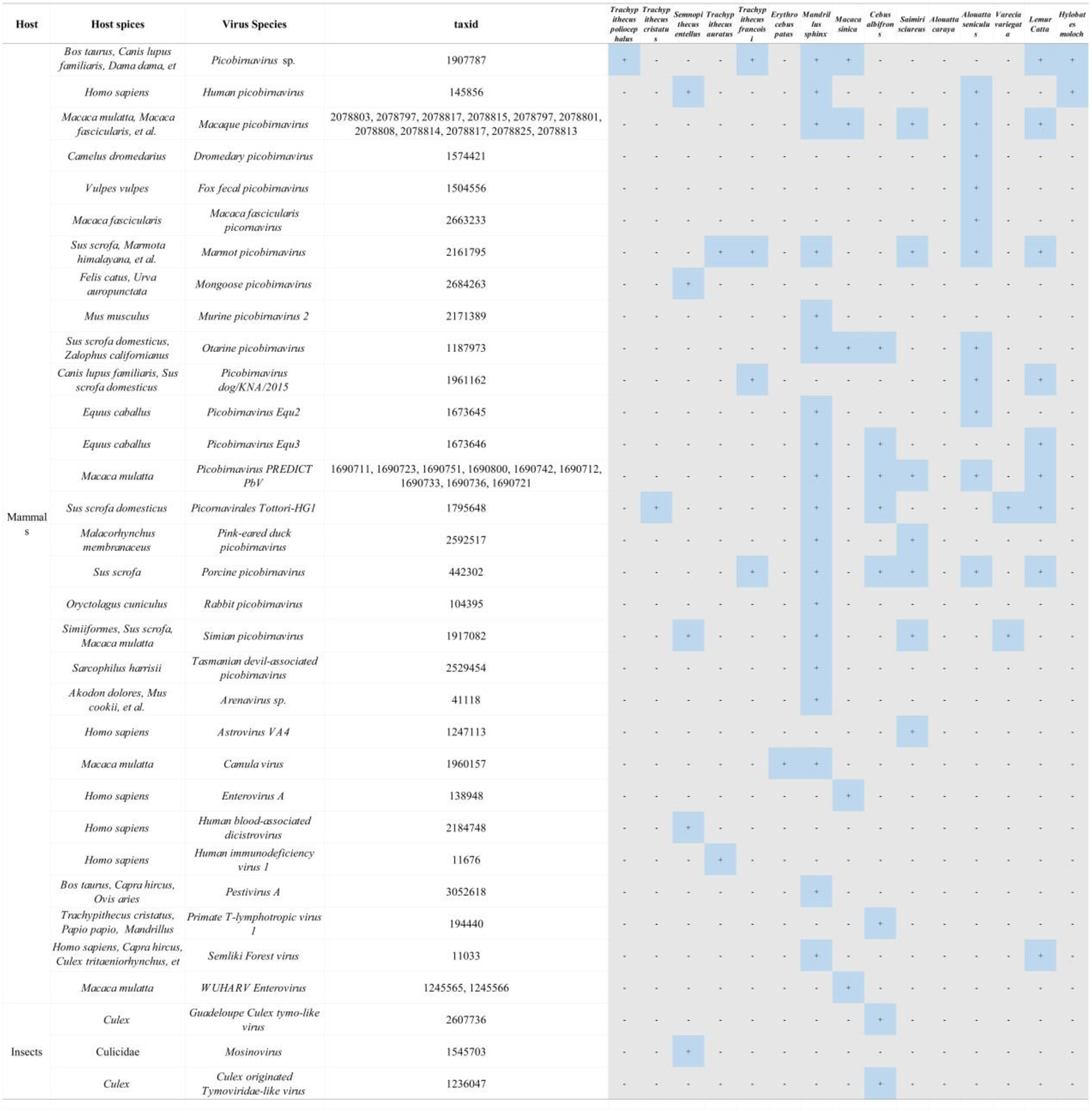
Potential pathogens and their host types counted in 15 samples.

### 2.5 Phylogenetic analysis of viruses with zoonotic potential

Some of the reads found in the 15 NHP fecal samples have sequence homology with the putative pathogenic viruses. Below, we describe these viruses in detail.

#### 2.5.1 *Human immunodeficiency virus 1* (HIV-1)

HIV-1 in our samples was found only in Javan langur. One sequence from Javan langur is used as query sequence. NCBI blast aligned sequences exhibited 90.36%-99.39% sequence identity (E-value = 0) at accession lengths of 607bp-9,781bp. Combined with other public isolates of HIV-1 downloaded from Genebank, a viral ML tree containing 129 sequences was constructed (bootstrap = 1000; best BIC model = TVM+F+R5) and colored by evolutionary branches. Phylogenetic analysis clustered evolutionary branches into five categories (Figure 4a).

**Figure 4.**
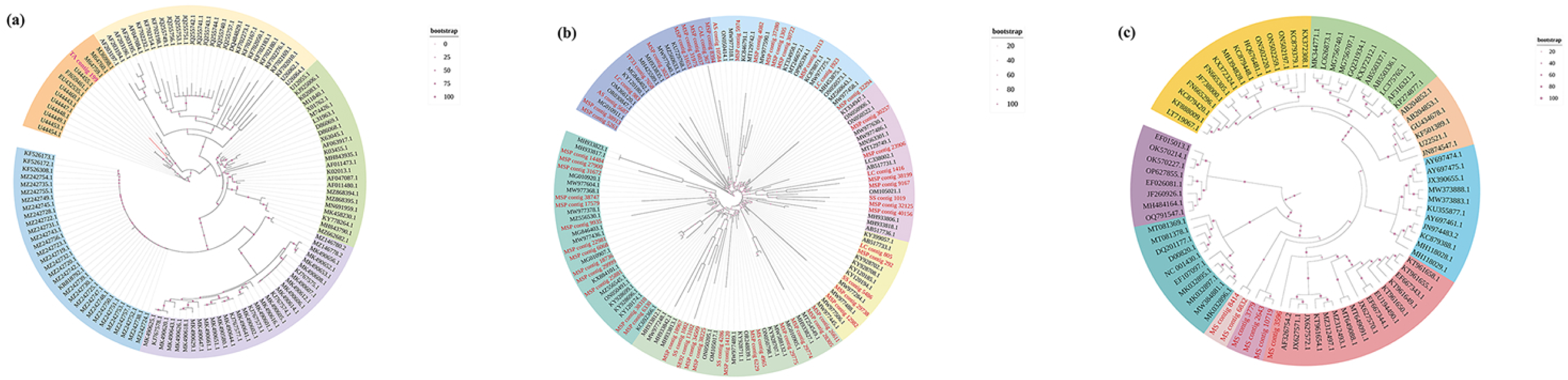
Phylogenetic tree of pathogenic RNA viruses, including the viruses identified in this study and related representative viruses. (a) Phylogenetic relationship of HIV-1 Nef region based on maximum likelihood method. bootstrap value is 1000, Best BIC model is TVM+F+R5; Which is on the same branch as the sequence obtained in our study is the human clinical HIV-1 Nef transcript (U44455.1) detected by the University of Washington in 1996. Sequences U44454.1, U44453.1, U44449.1, U44448.1, U44447.1, U44443.1, U44460.1 clustered on the orange branch are the same source as U44455.1. EU432535.1, FJ659461.1 from HIV-1 Nef transcripts detected in Venezuela in 2009(Rangel et al.,2009). M36998.1, M64759.1, M64760.1 from HIV-1 Long Terminal Repeat (LTR) Sequence isolated in 1990 by the Laboratory of Molecular Virology, New York, USA. (b) Phylogenetic relationship of PBV based on maximum likelihood method. bootstrap value is 1000, Best BIC model is GTR+F+R5; Sequences with similar evolutionary relationships to our reads came from respectively: Fecal samples from Cameroonians with diarrhea in 2019 (MH933827.1, MH933806.1, MH933818.1, MH933835.1); Human fecal samples from Kolkata, India, 2011 (AB517733.1, AB517736.1, AB517731.1); Pig feces samples from Manhattan, Kansas, USA in 2022 (MG010911.1); Domestic swine feces in North Carolina, USA, 2018 (MW977488.1, MW977506.1, MW977458.1, MW977590.1, MW977318.1); Fecal samples from pigs with diarrhea in August 2015 in the St. Kitts Island of the Caribbean (MT254549.1, MT246672.1); Fecal samples from domestic pigs in KwaZulu-Natal Province, South Africa, in March 2021 (OM105017.1, OM105021.1); Fecal samples from domestic pigs in Hunan, China, from November 2011 to June 2012 (KC846791.1); Pig feces samples (MZ556545.1, MZ556530.1) and cattle feces samples (ON050205.1, ON050798.1, ON050522.1, ON050414.1) from September 2017 in Hubei, China; Cattle feces sample from Bwindi, Uganda, January 2015 (OP905394.1); Cattle fecal samples from Hong Kong, China, from 2006 to 2013 (KY120180.1); Rhesus monkey feces from Bangladesh in 2013 (KT334958.1); Feces from SIV-infected rhesus monkeys in a 2015 study in Missouri, USA (MG010907.1, MG010905.1); Agile wallaby feces from Queensland, Australia (OR030847.1); California sea lion feces from zoos in Hong Kong, China (KU729768.1); Rabbit samples from the Australian Capital Territory (MT129749.1, MT129742.1); Red fox fecal samples collected in 2012 from southern Flevoland, The Netherlands (KC692366.1, KC878871.1); Himalay marmost feces collected from the Yushu Tibetan Autonomous Prefecture plateau, Qinghai, China, June-August 2013 (KY928707.1, KY928702.1); Lesser short-tailed bat fecal samples from New Zealand in 2020; Invertebrate samples from Jingmen, Hubei, China (KX884101.1) (c) Phylogenetic relationship of EV based on maximum likelihood method. bootstrap value is 1000 and Best BIC model is GTR+F+R10; The sequence MS contig8414 obtained in this study is in the same branch as MS contig6832; MS contig3779, MS contig2643 and MS contig10719 are in the same branch. Sequences clustered with MS contig3596 were derived from respectively: Rectal swab of Crab-eating Macaque submitted in a study in Virginia, USA (AF326754.2, enterovirus A122); Feces of rhesus monkeys suffering from SIV in a 2012 study at the University of Washington (JX627570.1: WUHARV Enterovirus, 1JX627571.1: WUHARV Enterovirus 2, JX627572.1: WUHARV Enterovirus 3); Diarrheal fecal samples from rhesus monkeys collected at the University of California, San Francisco in 2014 (KT961654.1: enterovirus A122; KT961658.1, KT961655.1, KT961655.1: enterovirus A124; KT961649.1: Simian enterovirus A92); Captive rhesus monkey feces collected in 2008 by the Centers for Disease Control and Prevention in Atlanta (EF667343.1: enterovirus A124, EF667344.1: Enterovirus A92); Feces of rhesus and pig-tailed macaque (Macaca nemestrina)collected in 2020 in Yunnan, China (MT649091.1, MT649088.1: enterovirus A122); Feces of rhesus monkeys with chronic diarrhea in Missouri in 2016 (MZ312494.1: Enterovirus J, MZ312497.1: enterovirus A122); Feces of rhesus monkeys reared at the Yerkes National Primate Research Center Field Station, USA, February-April 1999 (EU194490.1: Enterovirus A92)

#### 2.5.2 *Picobirnavirus* (PBV)

PBV was detected in our samples from several species, including mandrill, squirrel monkey, northern plains gray langur, toque macaque, ring-tailed lemur, colombian red howler, white-fronted capuchin, and Francois’ langur. 59 sequences were used as query sequences. NCBI blast aligned sequences showed 70.48%-98.18% sequence identity (E-value < 1e-5) at accession length of 337bp-2285bp. Combined with other public isolates of PBV downloaded from Genebank, a viral ML tree containing 138 sequences was constructed (bootstrap = 1000; best BIC model = GTR+F+R5) and colored by evolutionary branches (Figure 4b). Phylogenetic analysis clustered evolutionary branches into seven categories and phylogenetic relationships showed high diversity.

#### 2.5.3 *Enterovirus* (EV)

EV in our samples was found only in toque macaques (*Macaca sinica*). five sequences were used as query sequences. NCBI blast aligned sequences showed 95.79%-72.28% sequence identity (E-value < 1e-5) at the accession lengths of 7,586bp-381bp. Combined with other public isolates of EV downloaded from Genebank, a viral ML tree containing 83 sequences was constructed (bootstrap = 1000; best BIC model = GTR+F+R10) and colored by evolutionary branching (Figure 4c). Phylogenetic analysis clustered evolutionary branches into nine categories.

## 3 Discussion

### 3.1 Captive NHPs with lower total viral numbers carry food-borne plant viruses and potential zoonotic pathogens

In this study, RNA meta-virome sequencing was used to obtain a total of 3,188 known viral sequences from 15 captive NHPs, covering 376 virus species and 33 viruses that could potentially infect vertebrates. Due to our monkeys being healthy individuals, the number of viral sequences obtained was apparently lower compared to individuals of monkeys with diarrhea (27). The much lower number of viruses compared to wild or free-range animals suggests that captive conditions are much more hygienic than wild environments (28, 29). In contrast to other studies reported from zoos, farms, or animal management organizations (30–32), our study did not identify significant zoonotic pathogens harmful to humans in captive NHPs. Given the lack of current research on NHP viruses outside of the genus *Macaca* (26), it remains debatable whether this result is due to the species’ inability to carry zoonotic pathogens themselves, or whether it is due to the circumstances of good captivity. The total number of viral sequences of hosts whose evolutionary relationships are on the same branch may have large differences, for example, colombian red howler (96,824 reads) and black howler (30,539 reads) (Table 1). Using the Pearson product-moment correlation coefficient test, we found that there was no significant correlation between the average evolutionary distance of the species and the total number of viral reads (*p*-value = 0.9874). And the sample correlation coefficient was very close to 0 (r = 0.004), indicating that the evolutionary distance of the hosts and the total number of their viral reads have almost no linear correlation. The more likely reason for the differences in virus numbers is the different diets and rearing standards of the enclosures. The 95% confidence interval (-0.509 - 0.516) also contains 0, further supporting this conclusion.

In addition, most of the viruses are plant viruses, such as *Hibiscus latent Fort Pierce virus* (HLFPV) and *Hibiscus latent Singapore virus* (HLSV), which are most prevalent in langurs and ring-tailed lemur. They take several species of plants of the Malvaceae as natural hosts, are pathogens of hibiscus leaf crinkle disease (33), and can usually be infected/carried by a wide range of plant leaves. This is consistent with the dietary data we observed, as langurs eat more leaves and the natural host of the virus in their diet, Hibiscus (*Hibiscus rosa-sinensis*), was recorded. However the ring-tailed lemur’s main diet is fruit and doesn’t include any leaves. This may be due to the monkeys’ daily diets that were rationed and distributed on time, and the virus was spread temporarily to the ring-tailed lemur’s food by leaves during the transportation and distribution process.

Screening of samples for pathogenic threats in captive NHP detected several potentially risky viruses including HIV, PBV and EV. Mosquito viruses such as MVV, CYV and EXV were also detected from our samples. These viruses present in the NHP can affect public health safety and pose a potential threat to human health. Given that South China is a subtropical, densely populated and well-populated area that creates favorable conditions for the flourishing of plant and animal insects and viruses, and is a prone area for zoonotic diseases, these zoonotic viruses that we found in these caged NHPs need further attention.

### 3.2 Javan langur may have the ability to be infected with HIV-1

Although HIV-1 RdRp sequences were not found, our study did detect transcripts associated with HIV-1. Specifically, we identified a sequence annotated as an HIV-1 transcript in a sample of Javan Langur, which has an evolutionary similarity to clinical HIV-1 *Nef* transcripts detected in AIDS patients at the University of Washington, USA, in 1996 (34) and Venezuela in 2009 (35). Nef is a small myristoylated protein of 27-35 kDa encoding primate lentiviruses (HIV-1, HIV-2 and SIV) (36, 37). In hosts infected with pathogens, expression of the Nef protein significantly promotes viral replication and increases viral load to rapid onset of disease (38, 39). Individuals infected with HIV-1 encoding a defective *Nef* gene do not develop AIDS for decades (36, 37). Nef is thus thought to be a key factor in the pathogenesis of AIDS. Although the HIV-1 RdRp was not detected by us, based on the fact that Nef is abundantly expressed early in the HIV-1 viral replication cycle (40), it may be the case that the viral load is very low and has failed to be detected for the time being. And, because sequencing data came from animals fecal samples rather than blood samples, it also reduces the likelihood that potentially low-abundance HIV-1 viruses will be detected by us.

The transmission of HIV-1 does not include fecal, unlike some other extremely harmful infectious diseases, such as SARS (41) and MERS (42). In this study, we detected the Nef protein, which is expressed early in HIV-1 replication, in fecal samples. Based on this finding, we believe that although fecal transmission is extremely unlikely in HIV-1 transmission, prediction by excretion of feces may be a possible approach; however, since HIV-1 RdRp was not measured in fecal samples, this possibility needs to be fully and carefully evaluated.

Another interesting perspective is that previous studies have suggested that only pig-tailed macaques in Old World monkeys have been shown to be infected with HIV-1 and develop AIDS, allowing them to be used as an animal model for AIDS (43–45). However, our study points to the possibility of HIV-1 infection in Javan langurs. Although viral infections may be a transient viral spillover phenomenon in the host and pathogens are eventually lost from the host without repeated reintroduction through cross-species transmission (46). Still, the finding implies the possibility of entirely new HIV-1 model animals being developed, which provides new insights into the lack of NHP animal models of AIDS capable of HIV-1 infection.

### 3.3 PBVs with great zoonotic potential are widespread in NHPs

PBV represented the highest abundance of all vertebrate viruses detected in our samples. PBV was present in faecal samples from hosts mandrill, squirrel monkey, northern plains gray langur, toque macaque, ring-tailed lemur, colombian red howler, white-fronted capuchin, and Francois’langur(*Trachypithecus francoisi*). We found that the detected PBVs of the NHP clustered with diverse vertebrate-associated gene clusters and did not form separate clusters strictly by host species, suggesting a high degree of diversity of PBVs of the NHP. This result is consistent with previous studies on other animal hosts, such as rabbits (47), horses, pigs, cows (48), and monkeys (49). PBV sequences from various animal species have been found to be widely distributed throughout the phylogenetic tree, with a high degree of diversity both within and between host species (48). And the identified PBV phylogenies suggest that these viruses found in humans and other animals are genetically related (50–52). As we found NHP (mandrill, squirrel monkey, Northern Plains Gray Langur) PBVs clustered in the same taxa as African Cameroonian human intestinal PBVs, suggesting that humans and NHPs are shared viral hosts. These NHP hosts carrying PBV have no apparent taxonomic consistency. Similarly, past molecular studies of PBVs have shown them to be poorly coherent with host taxonomy (53), highlighting the lack of understanding of PBV dynamics.

Most previous studies have considered PBV to be an opportunistic pathogen (54) and proposed its association with gastroenteritis and acute watery diarrhoea in humans (55–57). In NHP-related studies, using a variety of virus-specific reagents and methods, diarrhoeic rhesus monkey faeces from monkey farms in China and the Yerkes National Primate Research Center (Yerkes, Georgia) have been shown to contain PBV (20); diarrhoeic crab-eating macaques and pig-tailed macaques have also been described as infected with PBV (57). It has also been suggested that PBV lacks conclusive detection in solid tissues and does not necessarily cause diarrhoea in vertebrates (58). Instead, they are more likely to infect prokaryotes due to the binding sites they possess (59). This interaction between PBV and prokaryotes may further explain the evolution and genetic diversity of PBV.

Woo et al. observed that the evolutionary mechanism of PBV may be similar to that of other segmented RNA viruses, such as the genome reassortment evolution of rotaviruses (48). This evolutionary mechanism has resulted in rotavirus high diversity, emergence of pathogenic strains and outbreaks of infectious diseases (60, 61). Consequently, PBVs with this evolutionary mechanism also have a high degree of genetic diversity, which likewise creates the possibility that just some of the many genotypes transmitted have pathogenic potential. We need to be bolder in our focus on PBV, a widespread opportunistic zoonotic pathogen that can cause diarrhoea, with a simple route of transmission and easy access to humans. The results of our data further confirm previous reports of high genetic diversity in PBV. In a study of PBV in monkeys, a ratio of Ka/Ks < 0.05 was observed for the protein-coding genes of PBVs, implying that the virus evolved stably in monkeys (49). Therefore further screening of PBV viruses of the NHP to explore the mechanisms of PBV diversity and evolution, and whether there is an association between them and NHP diarrhoea will shed more light on the epidemiological potential threat to human health from animal viruses (3).

### 3.4 EV was found in seemingly healthy monkey individuals

EV can infect a wide range of mammals including humans and has strong zoonotic characteristics (62). However, although EV may transmit between NHP and humans to each other (63), it is not usually associated with disease in monkeys (64). Most of the serotypes that have been identified in the past are human pathogens. Several common serotypes isolated from humans, *enterovirus A71*, *coxsackievirus A16*, and *coxsackievirus A6*, have been instrumental in HFMD outbreaks in the Asia-Pacific region (65, 66). Currently, several serotypes EV-A122, A123, A124 and A125 have also been isolated from apes (67).

The EVs obtained by sequencing in our study were found to have a similar evolutionary relationship with *Enterovirus A122*, *Enterovirus A124*, *Enterovirus A92*, *Simian enterovirus A92*, *Enterovirus J*, *WUHARV Enterovirus1*, *WUHARV Enterovirus2*, and *WUHARV Enterovirus3* found in other NHPs have similar evolutionary relationships and form a single lineage in the phylogenetic tree. Moreover, most of the NHPs that detected EV in previous studies were species of the *Cercopithecinae* subfamily in old world monkeys, such as crab-eating macaques (68), rhesus monkeys (21, 27, 69, 70), Pig-tailed Macaques (71), baboons (72, 73), and African green monkeys (74). This is in accordance with our findings, as we detected EV only in the toque macaques sample. These results suggest that hosts of the *Cercopithecinae* appear to be more favoured by EV, while the virus is most likely to be shifted to humans through these host species. These newly identified NHP EVs have rarely been reported (75), however, their potential for spillover to humans is very significant due to the ability to be spread through the faecal-oral or respiratory tracts.

## 4 Conclusion

Overall, this study suggests that captive NHP harbour zoonotic pathogens or viruses with an evolutionary potential to cause disease. Although the majority of viruses in captive NHPs were associated with their plant food, *Picobirnavirus* (PBV), *Enterovirus* (EV), Nef protein transcripts of *human immunodeficiency virus 1* (HIV-1), and insect viruses, which are related to human and other mammalian viruses on the phylogenetic tree, were also detected. Remarkably, these seemingly healthy captive NHPs can also carry infectious pathogens. This study highlights the risk of zoonotic disease carriage and transmission by NHP and complements the information on viruses harbored by NHP species for which there is insufficient clinical surveillance.

## 5 Materials and methods

### 5.1 Samples collection

During November-December 2019, we sampled at a zoo in southeastern China. A total of 45 samples from 15 adult male primate individuals were obtained. These individuals include: Javan Silvery Gibbons (n = 1), Black Howler (n = 1), Colombian Red Howler (n = 1), White-Fronted Capuchin (n = 1), Squirrel Monkey (n = 1), Ring-Tailed Lemur (n = 1), Black-and-White Ruffed Lemur (n = 1), Patas Monkey (n = 1), Toque Macaques (n = 1), Mandrill (n = 1), Francois’Langur (n = 1), Cat Ba Langur (n = 1), Silvered Langur (n = 1), Javan Langur (n = 1), Northern Plains Gray Langur (n = 1). Host species information was provided by the zoo. Each species contained 1 focal individual, and individual focal animals were observed continuously and feeding data were recorded.

Fecal samples for high-throughput sequencing of the meta-virome were collected and stored in 2 mL centrifuge tubes using sterile cotton swabs and sterile toothpicks immediately after defecation on the same day as the focal individual, snap-frozen in liquid nitrogen, and then transferred to a -80℃ refrigerator for storage. Three faecal samples were collected from each individual and all faecal samples were labelled with information on the time of collection, species and specification.

### 5.2 Library preparation, quality control and reads assembly

As our samples came from a clean captive environment with low levels of RNA viruses, samples from each pool were pretreated by ultracentrifugation method (MAGEN, Guangzhou, China) to achieve high purity RNA extraction. Sample pre-treatment, viral nucleic acid extraction methods are described in Appendix 1. Quality control and reads assembly methods are described in Appendix 2.

### 5.3 Virus identification, abundance statistics and gene prediction

Viral sequences were identified using the Denovo method, details of the methodology and exclusion of false positives are given in Appendix 3. Species annotations were made based on BLAST (v2.9.0+) comparisons of viral contigs to the NT database, selecting best hit comparisons with e ≤ 1e-5, and no comparisons are denoted by NA. BWA software (v0.7.17, parameter: mem-k 30) compares clean reads with each identified virus contigs, filters out reads with comparison length < 80%, then calculates the distribution of virus reads based on the annotation results of the virus contigs, and finally calculates the RPKM for each virus. MetaGeneMark software (76) (v3.38) was used for gene prediction of virus contigs, filtering sequences with gene nucleic acid lengths less than 300bp.

### 5.4 Phylogenetic analysis of pathogens

For phylogenetic analyses, pathogen sequences screened in 15 samples were used as bait for comparison at NCBI (value = 1e-5) and representative viral genomes or gene sequences were downloaded from GenBank. IQ-tree (77) was used to determine the best model as well as to construct a maximum likelihood (54) tree (bootstrap=1000). Visualisation of phylogenetic trees was performed using Interactive Tree of Life (iTOL, https://itol.embl.de/) (78).

### 5.5 Statistical analysis

The evolutionary tree of the host species was constructed using timetree (https://timetree.org) to obtain the Newick file used to calculate the evolutionary distance. The R language ‘ape’ package was used to extract a Patristic distances between host species, where the distance between each pair of species reflects the length of their branches in the evolutionary tree. To ensure consistency between the patristic evolutionary distance matrix and the viral reads data, we performed a strict correspondence matching between the two datasets, and the average evolutionary distance for each species was calculated for comparison with the corresponding species viral reads numbers. The cor function in R was used to test the Pearson correlation between these two data sets. The significance level for all statistical tests was set at 0.05.

## Authors contribution

**Yujie Yan:** Writing--original draft; data curation; formal analysis; visualization. **Yuhang Li:** Data curation; investigation; validation. **Linshan Yang:** investigation; validation. **Haojie Wu:** investigation; validation. **Fan Wu:** Investigation. **Hongli Chang:** Supervision. **Zhengfeng Hu:** Investigation. **Shujun He:** Investigation. **Yi Ren:** Investigation. **Lifeng Zhu:** Writing--review and editing; methodology. **Baoguo Li:** Writing--review and editing. **Songtao Guo:** Writing--review and editing; conceptualization.

## Acknowledgement

This work was supported financially by the Major International Joint Research Program of Natural Science Foundation of China under Grant (32220103002); National Natural Science Foundation of General Project under Grant (32370534); Natural Science Foundation of China under Grant (32371563); and Innovation Support Plan of Shaanxi Province under Grant S2021-ZC GHID-0013.

## Conflict of interest disclosure

The authors declare no conflict of interest.

## Ethics approval statement

A total of fresh feces of captive NHPs were collected with the permission of the authorities of a zoo in southern China. All procedures were approved by the Ethics Committee for Experimental Animal Management and Welfare of the Northwest university.

